# Similar distribution of T-cell subsets in Crohn’s disease and in diverticulitis provide evidence against a primary causal role for these cells in Crohn’s

**DOI:** 10.1101/861807

**Authors:** Ayse U Akarca, Peter Ellery, Anthony W Segal, Teresa Marafioti

## Abstract

**Background and Aims:** T lymphocytes are found in abnormally large numbers in the bowel in Crohn’s disease. This has led to the assumption by some that these cells play a causal role in the pathogenesis of what has been labelled an autoimmune disease. An alternative explanation for their presence is that, as part of the adaptive immune system, the accumulation of these cells is not a primary phenomenon, but is a secondary adaptive immune response to faecal material in the bowel wall. To distinguish between these two processes we compared the T-cell repertoire in the bowel in Crohn’s with that in diverticulitis, where the primary pathology is mechanical, with a subsequent immune response to the accumulated faecal material.

**Methods:** Six cases of Crohn’s disease and six patients with diverticulitis were studied. Dewaxed sections of bowel were stained with Anti-CD4, Anti-CD8, Anti-FOXP3 and Anti-CD25 to identify cytotoxic T-cells, NK-Tcells; T-helper and T-reg T-cells.

**Results:** No differences were found in the distribution of the different T-cell markers in either the mucosa or in areas of inflammation in the two conditions.

**Conclusion:** The accumulation of T-lymphocytes in the bowel in Crohn’s disease is likely to be a sign of an adaptive immune response to faecal material within the bowel rather than an indication of a primary causal immune attack on the bowel that produces the disease.

## Introduction

The clinical picture of an inflamed bowel containing large numbers of macrophages and T-cells^12^ has led to the belief that Crohn’s disease is a T-cell dependent autoimmune disease^3^. This concept has been supported by experiments in mice ^45^ which have sparked a large body of work into the role of regulatory T cells in the pathogenesis of IBD.

More recently, interest in the role of T-cells in the pathogenesis of Crohn’s disease has switched from T-cells in general to different subsets of T-cells, including T helper (Th)1 cells, Th17 cells, and regulatory T-cells^2^.

However, over the past decade, multiple groups have failed to find abnormalities in these cells in the intestines or blood of patients with IBD^3^.

Diverticulitis results from the herniation of a pouch of mucous membrane, including muscularis mucosae, through and beyond the circular muscle layers of the bowel wall into the pericolic fat. Faecal material trapped in these sacs can penetrate locally and cause sepsis and inflammation, with an inflammatory response indistinguishable from that in CD^6^ with an intense infiltration by lymphocytes.

It is possible that different T-cell subsets infiltrate the bowel wall in Crohn’s disease and diverticulitis. Were this to be the case the different T-cells could play different pathogenic roles.

We have undertaken this investigation to compare and contrast the distribution of T-cell subsets in Crohn’s disease and in diverticulitis.

## Materials and Methods

### Patients

Six cases of Crohn’s disease and six patients with diverticulitis were studied. All cases were obtained from the routine diagnostic archives of University College Hospital, London, UK. Each case was diagnosed histologically by an experienced histopathologist, and on the clinical information, and reviewed by TM and AWS.

### Immunohistochemistry

Sections were cut from paraffin embedded blocks, transferred onto poly-l-lysine–coated slides and dewaxed. Single immunohistochemistry was carried out using the automated platforms BenchMark Ultra (Ventana/Roche) and the Bond-III Autostainer (Leica Microsystems), and multiplex immunohistochemistry, were performed using protocols described previously^7^. Co-expression of nuclear and cytoplasmic or membranous proteins was clearly detected, as the colour of the chromogens remained distinct. Images were scanned using the NanoZoomer Digital Pathology System C9600 (Hamamatsu).

Mouse monoclonal primary antibodies were used to identify cytotoxic T-cells (Granzyme B, CD8); NK-Tcells ((granlysin); T-helper (CD4) and T-reg (CD4, FOXP3), (FOXP3, CD25) T-cells. All the antibodies were from Leica Microsystems, UK, apart from Anti-FOXP3 which was a gift of Dr G Roncador, CNIO, Madrid, Spain. Clones used were: Anti-CD4, 4B12; Anti-CD8, 4B11; Anti-FOXP3, 236A/E7; Anti-CD25, AC9. Human reactive tonsil was used as the positive control.

### Analysis

For each patient an obvious area of inflammation in the bowel wall, or an area of mucosa, was randomly selected. Five representative areas (0.25 mm^2^) were selected within each area of bowel inflammation or mucosa.

Single positive CD4, CD8 cells and double positive CD4 Foxp3, CD25 CD4 cell were quantified in QuPath software. The average cell count per 1.25 mm^2^ were calculated separately for areas of inflammation, and mucosa.

## Statistical analysis

Statistical analyses were performed with GraphPad Prism 7 (GraphPad Software); p values were calculated using Student’s t test to compare unpaired groups (ns = p > 0.05).

## Results

No significant differences were found in the absolute numbers, or the relative distribution, of the T-cell subsets in inflamed bowel or mucosa from patients with Crohn’s disease or with diverticulitis (Figures).

**Figure 1.**
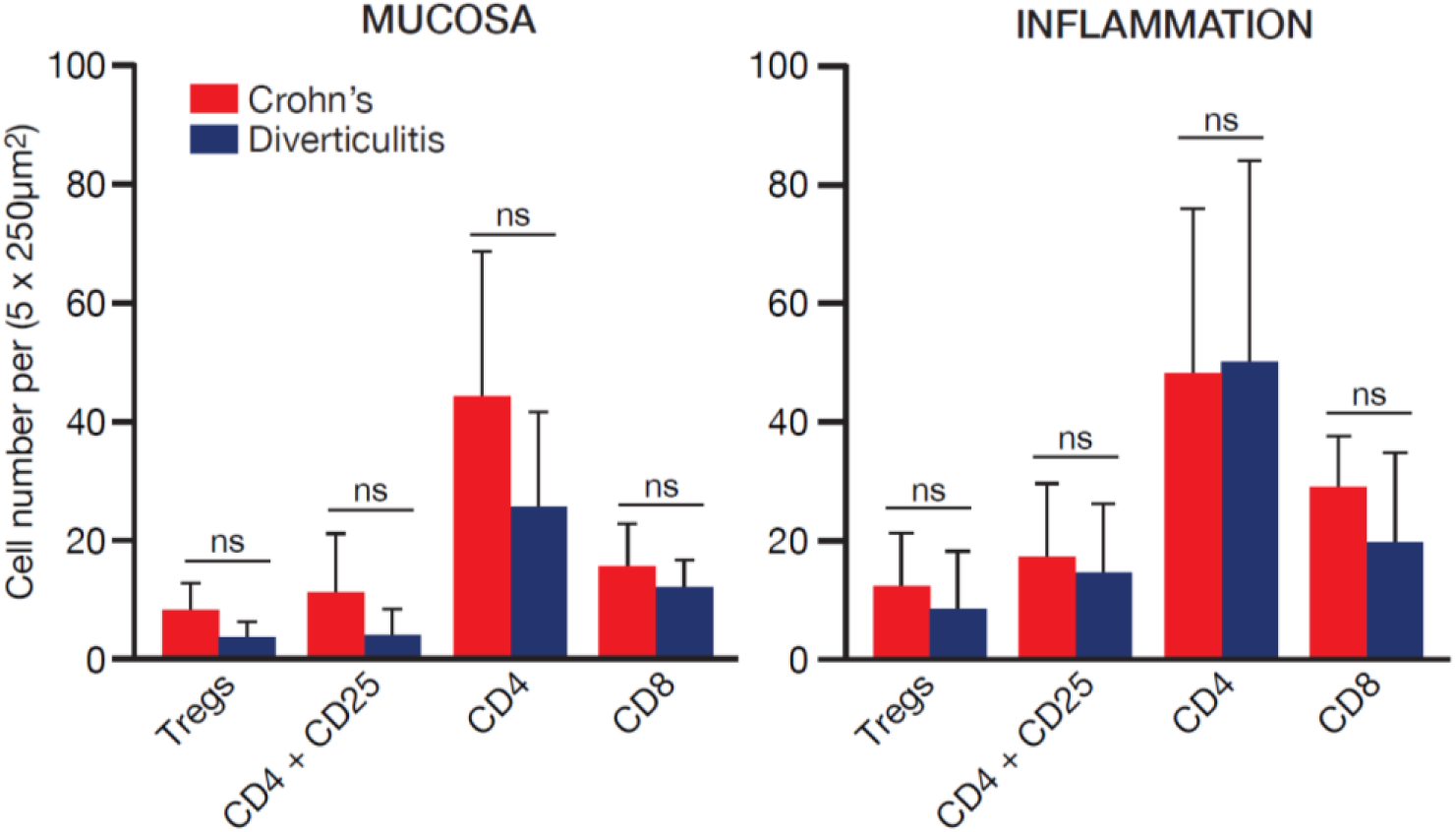
Numbers of the different T-cell subsets in the mucosa and in areas of inflammation within the bowel wall (mean + SD). Counts were made in five randomly selected areas of 250μm2 in each of the six patients with Crohn’s disease or with diverticulitis. No significant differences were found in the T-cell subsets in the two diseases (ns = p > 0.05) 1 and 2).

**Figure 2.**
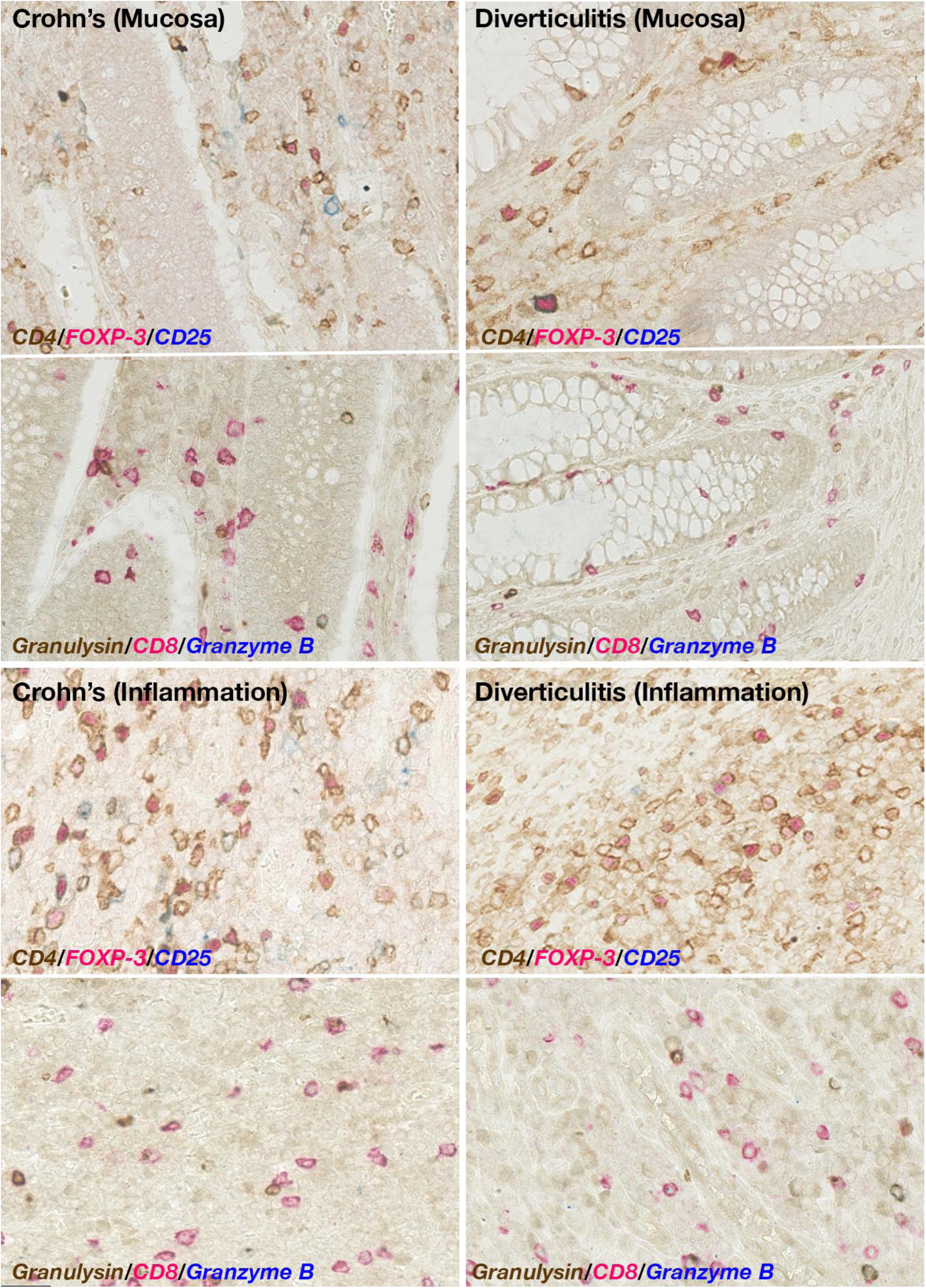
Typical examples of multiplex immunohistochemistry staining patterns of sections of mucosa and inflamed bowel from patients with Crohn’s disease and diverticulitis. The distribution of T- cell subsets is shown in areas of the mucosa and of inflammation.

## Discussion

The underlying pathological process in Crohn’s disease has been demonstrated to be a weak acute inflammatory response. The consequences of this are the depressed recruitment of neutrophils to lesions, which in the bowel results in the failure of the clearance of faecal material that gains access to the tissues^8^. This faecal material is then walled off, leading to a chronic granulomatous inflammation and an adaptive immune response. Diverticulitis also leads to the penetration of faecal material into the wall of the bowel. Both conditions result in a similar accumulation of T-cells in the affected tissues despite very different routes to the development of the inflammation.

It is difficult to conclude other than the accumulation of T-cells in the affected bowel in Crohn’s disease is an adaptive response to the bacteria that have gained access to the tissues, rather than their presence representing evidence of a causal T-cell mediated immunological attack.

These findings have significance for the direction that future therapeutic strategies should take.

## Acknowledgements

Approval for this study was obtained from the National Research Ethics Service, Research Ethics Committee 4 (REC Reference number 09/H0715/64).

## Author Contributions

AWS and TM planned the experiments, AA and PE conducted the experiments, AA and TM analysed the immunostaining, AWS and TM wrote the manuscript.

